# Dual-channel graph learning reveals similarity and complementarity in protein-protein interaction networks

**DOI:** 10.64898/2026.01.26.701665

**Authors:** Tao Tang, Taiguang Shen, Weizhuo Li, Yangyang Chen, Sisi Yuan, Yuansheng Liu, Xinyu Yang, Xiao Luo

## Abstract

Protein-protein interactions (PPIs) are governed by two fundamental interfacial mechanisms: similarity-driven, often involving symmetric structural motifs, and complementarity-driven, arising from geometric and physicochemical matching between binding surfaces. Despite their biological significance, computational models have largely overlooked the coexistence and interplay of these twofold interaction modes. Here, we introduce DMG-PPI, a dual-channel graph neural network framework that jointly models similarity and complementarity in PPI networks, extending prior heterophilous GNN concepts to explicitly disentangle these dual interaction modes. The model consists of two parallel message-passing pathways: Alignment Message Passing (AMP), which aggregates signals from similar neighbors to capture symmetric interfaces, and Divergence Message Passing (DMP), which contrasts node features to extract complementary binding patterns. The signals captured by AMP and DMP are integrated via an adaptive fusion strategy within each block, and the outputs of blocks are aggregated using the MixHop framework to encode higher-order interaction patterns. DMG-PPI substantially outperforms state-of-the-art methods on classical benchmark datasets, achieving a 7.19% improvement in Micro-F1 over the second-best method. Additionally, the dual-channel framework provides interpretable insights into key binding residues by identifying interfacial mechanisms. Overall, DMG-PPI serves as a powerful tool that reveals the mechanisms behind accurate PPI predictions and facilitates downstream biological analysis.

**Author summary:** Proteins interact through multiple interfacial principles rather than a single uniform mechanism, reflecting diverse modes of molecular recognition between binding partners. Accurately modeling this heterogeneity remains a central challenge in protein–protein interaction (PPI) prediction. Existing computational approaches often overlook such complexity and treat all interactions as arising from a dominant pattern. To address this limitation, we present DMG-PPI, a dual-channel graph neural network framework designed to account for distinct interaction principles within a unified model. By modeling both similarity-based and complementarity-based interaction signals, DMG-PPI enables a more comprehensive representation of interaction patterns in PPI networks. Evaluations on widely used benchmark datasets demonstrate that our approach achieves robust and consistent improvements over existing methods. Beyond prediction accuracy, DMG-PPI provides biologically meaningful interpretability by highlighting key residues and revealing distinct interfacial interaction mechanisms, offering valuable insights for downstream structural and functional studies.

## Introduction

Protein–protein interactions (PPIs) constitute the structural and functional backbone of cellular systems, coordinating molecular processes ranging from signal transduction to metabolic regulation [1, 2]. PPI networks therefore provide a systems-level representation of how biological function emerges from molecular interactions and how their disruption drives disease. Mapping and modeling these networks is central to both fundamental biology and biomedical research, enabling insights into molecular organization [3], complex assembly [4], system-level cellular behavior [5],target prioritization [6], and therapeutic intervention [7]. Although experimental techniques have generated invaluable PPI data, their high cost, context dependence, and limited scalability necessitate computational approaches capable of predicting interactions at the network scale.

At a fundamental level, protein binding is governed by two non-mutually exclusive interaction modes. **Similarity-driven interactions** arise from shared evolutionary or biochemical features that favor compatible binding affinities [8, 9], whereas **complementarity-driven interactions** depend on precise geometric and physicochemical matching at binding interfaces [10, 11]. Experimental evidence supports the coexistence of these interaction modes: Yeast two-hybrid assays preferentially detect interactions between proteins with similar domain architectures [12, 13], whereas X-ray crystallography reveals complementary binding at atomic resolution [14]. This duality is not unique to PPIs but recurs across biological networks, including gene regulatory [15–17], signaling [18–20], and metabolic systems [21, 22], highlighting a general organizing principle of biological systems. This simplification reflects a deeper conceptual mismatch between biological interaction mechanisms and their computational abstraction. In particular, PPI networks cannot be faithfully represented by models that conflate similarity and complementarity into a single interaction logic. Consequently, the dual nature of PPI formation has not been systematically formulated as co-equal, first-class modeling principles in computational interactomics, limiting our ability to disentangle, interpret, and reason about distinct interaction modes at the network level.

Currently, the research in computational PPI prediction primarily addresses two key aspects: learning informative representations of individual protein from their sequences or structures [23, 24], and modeling the interaction patterns within PPI networks [25]. The former aspect primarily relies on physicochemical characteristics [26, 27] and residue-level encoding [28, 29] of protein sequences to generate protein embeddings. More recently, Gao et al. [30] introduced graph neural networks (GNNs) to model protein tertiary structures as graphs. Graph autoencoder and masked modeling was then applied to learn a robust representation of protein structure [31]. The latter aspect concentrates on modeling interactome using GNN. Graph Isomorphism Network (GIN) was among the first GNN architectures applied to PPI prediction [32]. Subsequent studies have demonstrated improvements in generalization capabilities through Graph Attention Network (GAT) [33] and anti-symmetric Graph Convolutional Network (GCN) [34], as well as incorporating GNNs into broader frameworks [35, 36].

While graph neural networks (GNNs) have become widely used for PPI prediction, most existing architectures primarily aggregate features from similar neighbors, effectively biasing them toward similarity-driven interactions and limiting their ability to capture complementarity-driven associations. Although prior work on heterophilous GNNs has introduced mechanisms to handle structurally or functionally dissimilar nodes, these approaches were not specifically designed to disentangle the co-occurrence of similarity- and complementarity-driven interactions inherent to PPI networks. This gap constrains both predictive accuracy and biological interpretability.

Here, we introduce DMG-PPI, a Dual Message Pathway Graph Neural Network that explicitly models the dual principles governing protein–protein interactions. DMG-PPI decomposes message passing into two parallel channels: Alignment Message Passing (AMP), which captures similarity-driven interactions, and Divergence Message Passing (DMP), which captures complementarity-driven interactions through contrasting features. These channels are processed in parallel and adaptively fused to generate multi-scale representations that preserve the distinct biological semantics of each interaction mode. By grounding network modeling in the physical principles of protein binding, DMG-PPI achieves superior predictive accuracy, robustness to noise, and enhanced interpretability, enabling inference of dominant binding modes and key interface features within PPI networks.

## Materials and methods

Homophily is a fundamental assumption in graph theory, referring to the tendency of links to form between similar nodes. This property is prevalent in many studies on graph structure, such as social network (Fig1a). Most previous studies of PPI networks have largely relied on this assumption, favoring GNN architectures with sum-based aggregation that naturally emphasize homophilous relationships. However, as illustrated in Fig 1b, biological evidence indicates that protein interactions often arise not only between structurally or functionally similar proteins but also between complementary counterparts, reflecting the coexistence of similarity- and complementarity-driven interactions.

**Fig 1:**
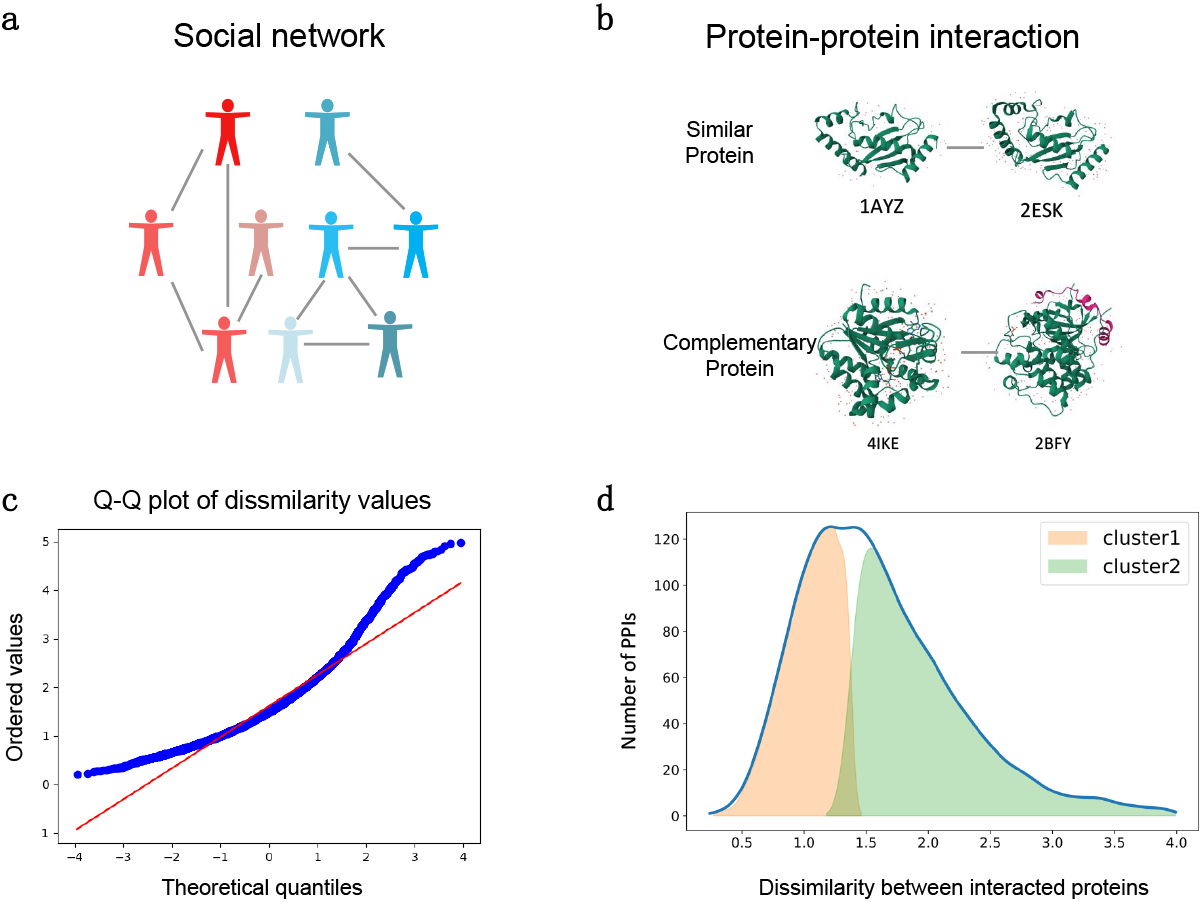
Protein–protein interactions (PPIs) can emerge through two principal mechanisms: similarity and complementarity. **a** In social networks, individuals with similar attributes tend to connect, a phenomenon termed homophily. **b** Analogously, in PPI network, both similar and complementary proteins tend to interact. **c** A Q–Q plot of dissimilarity values among interacting proteins corroborates this pattern: numerous points deviate substantially from the reference line, suggesting a departure from unimodality and supporting a mixture model. **d** Clustering analysis further categorizes the binding protein pairs into two groups exhibiting markedly different levels of structural dissimilarity. These groups correspond to interactions primarily driven by structural similarity and complementarity, respectively.

To systematically investigate protein interaction patterns, we quantified the similarity between interacting proteins as the euclidean distance between their embeddings derived from pre-trained geometric model [31]. A Q–Q plot of similarity scores for interacting pairs in SHS27K is shown in Fig 1c, where numerous points deviate substantially from the reference line, suggesting a mixed distribution of similarity among interacting proteins. Subsequent clustering analysis (Fig1d) also supports the coexistence of connection patterns in PPI networks: the distribution of similarity exhibits two distinct peaks, indicating the presence of two separable groups of protein interactions that likely correspond to similarity- and complementarity-driven relationships. Together, these results underscore the complex nature of PPI, in which both similarity-based and complementary interactions jointly contribute to network connectivity.

Here, we propose DMG-PPI for modeling the two-fold interaction patterns in PPI network. As depicted in Fig 2, our approach comprises four main modules: (1) construction of the physical protein graph, (2) a graph autoencoder for data preprocessing and generation of initial node embeddings, (3) a dual-channel GNN framework for capturing complex interaction patterns within the PPI network, and (4) a classifier for final prediction.

**Fig 2:**
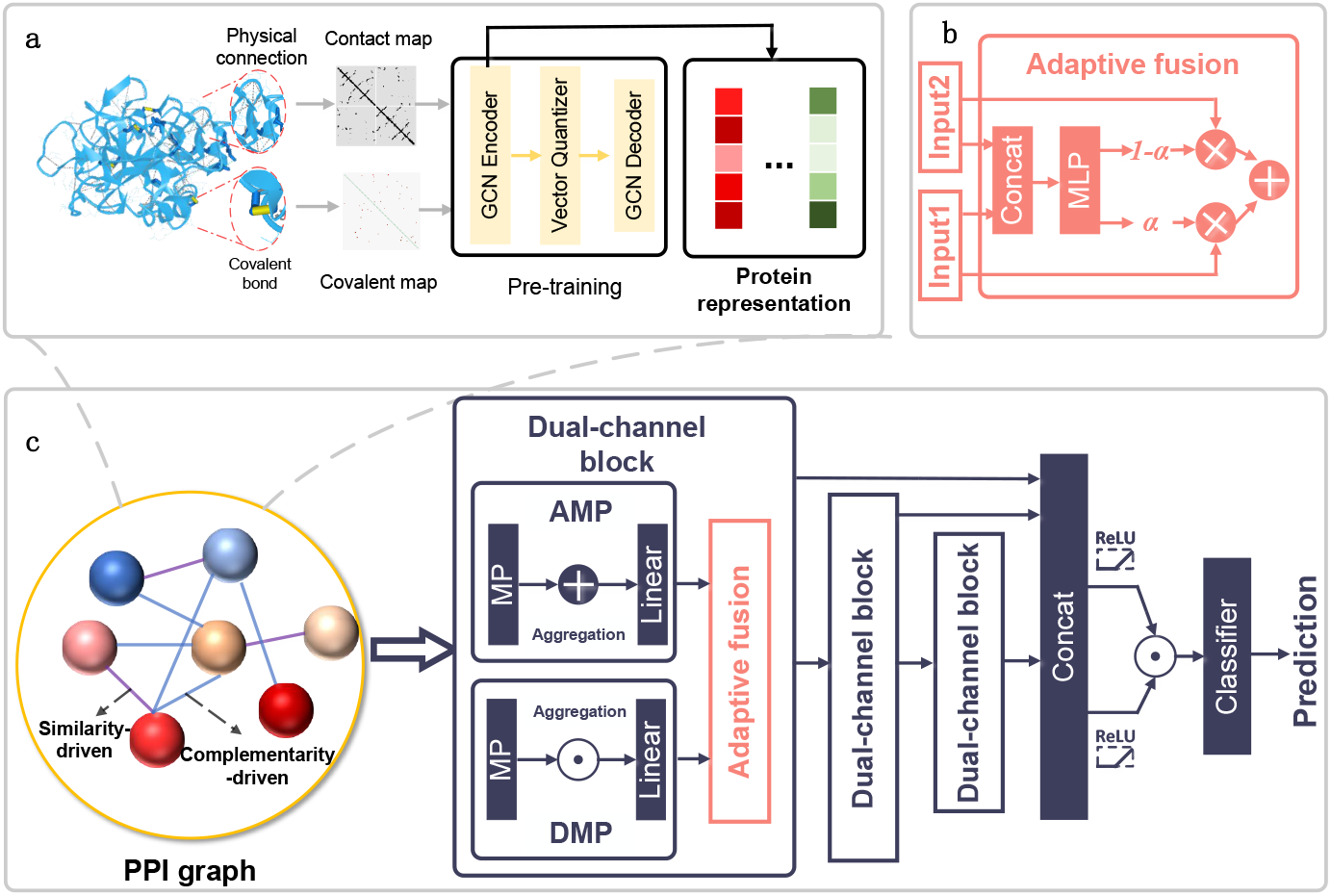
A schematic diagram of DMG-PPI. **a** The contact map and covalent map are processed by GNN autoencoder with a vector quantizer pre-trained on human interactome. The encoder output serves as the initial protein representations. **b** The adaptive fusion block of DMG-PPI fuses its inputs using a gating mechanism for context-aware integration. **c** Protein representations are recursively updated through dual-channel blocks, in which the input is processed in parallel by GCNs employing additive aggregation (AMP) and multiplicative aggregation (DMP). The outputs of each block are concatenated within a MixHop framework. Finally, the Hadamard product of these protein representations yields the PPI-level descriptor, which is then passed to a classifier for PPI prediction.

### Problem formulation

The PPI network is denoted as 𝒢 = (𝒫, ℐ), where 𝒫 = {*p*_1_, *p*_2_, …, *p*_*n*_} is the protein set, and ℐ = (*p*_*i*_, *p*_*j*_, *y*_*ij*_), the protein feature of *p*_*i*_ is defined as *x*_*i*_. The objective is to learn a model 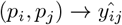 from training set.

### Feature extraction from intramolecular physical and chemical connections

We use residue-level graph to model the tertiary structure of each protein. Unlike the previous methods that only learn the spatial contact inside protein, our approach considers both the physical and chemical connections. For each protein, two residual-level protein graphs are constructed to model the physical and chemical connections respectively, both graphs consist of an adjacency matrix *A*_*p*_ ∈ {0, 1} ^*r×r*^ and a feature matrix *M*_*p*_ ∈ ℝ^*r×s*^, where *r* denotes the residue number of protein.

For chemical protein graph, the adjacency matrix is derived from covalent bonds between atoms; if there is a covalent bond between two atoms, then there is a connection between the corresponding residues. Covalent bonds are identified based on known chemical bonding patterns within amino acid residues and ligand molecules. For each pair of atoms, the connection is determined by the presence of a covalent bond as defined by standard residue chemistry or explicit bonding information in the structure data. The node feature is formed by three physicochemical properties: solvent accessible surface area (SASA), electrostatic potential surface (EPS), and pKa shifts. For the physical connection protein graph, a contact map is used as the adjacency matrix, in which the connection between residues is determined by the 3D coordinates of alpha carbon atom in residue. For each pair of residue, the connection exists if the distance between their alpha carbon atoms is smaller than 8Å. The node features are extracted by the type-dependent encoding of RDKit [37]. The physical and chemical protein graphs are processed with graph autoencoder to learn the embeddings of protein graph, we adapt the masked vector quantizer [31] for training process. The protein graph are processed by GNN encoder, GNN decoder and a vector quantizer:

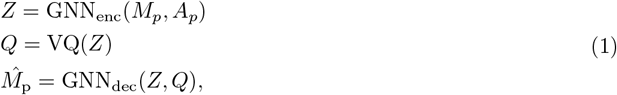

the masked modeling *m* is applied on output of vector quantizer:

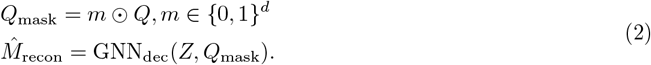

The graph autoencoder is pre-trained on SHS148K. The reconstruction loss is defined as the sum of three components: the mean squared error (MSE) between quantized representations *Q* and latent representation *Z*; the MSE between the original feature *M*_*p*_ and its reconstruction 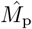; and the MSE between original feature *M*_*p*_ and masked reconstruction 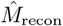.

The embedding of each protein *h*_*i*_ is obtained by concatenating the latent representations from the physical and chemical graphs, followed by a readout operation: *h*_*i*_ = READOUT(*Z*_phy_ ∥ *Z*_che_), ∥ denotes the concatenation operation.

### Dual-channel framework for learning interaction pattern

DMG-PPI employs two parallel channels to jointly model similarity- and complementarity-driven interactions simultaneously. Given a PPI graph 𝒢 = (𝒫,ℐ), along with the initial node embeddings **h** for each protein, DMG-PPI propagates information through a stack of dual-channel GNN blocks, 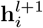 denotes the output of *l*th block. Within each block, protein representations are independently processed through two specialized message-passing pathways: AMP, designed to capture similarity-driven interactions, and DMP, tailored for complementarity-driven interactions. The outputs of AMP and DMP are formally defined as:

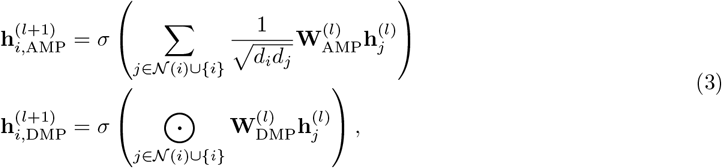

where 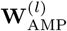 and 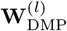 denote the learnable weight matrix of each channel, 𝒩 (*i*) denotes the neighborhoods of *p*_*i*_, *d*_*i*_ denotes the degree of the *i*th protein. AMP and DMP use summation and multiplicative message pathway respectively, to fit the biological characteristic of PPI network: summation aggregates features from neighbors and tends to smooth the representations, naturally reinforce similarity-driven PPI; multiplicative emphasize contrast and alignment between node features, aligns well with complementary-driven PPI.

A gating mechanism is introduced to adaptively balance the contribution of both channels. The gate coefficient *α* ∈ (0, 1) is obtained by linearly projecting the concatenated AMP and DMP embeddings:

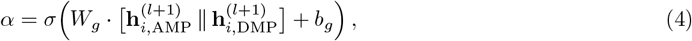

where *W*_*g*_ and *b*_*g*_ are learnable parameters, *σ* denotes an activation function that constrains *α* in range (0, 1). In this study, we employ sigmoid function. This coefficient adaptively regulates the relative contribution of each channel to the fused representation, as follows:

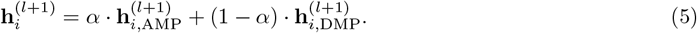

To capture multi-scale dependencies and alleviate the over-smoothing effects commonly observed in deep GNNs, we adopt a MixHop-based aggregation scheme for computing the final protein representation. In the MixHop framework, the embedding of each block is concentrated together, followed by linear transformation, batch normalization and activation function sequentially. The final node embedding is as following:

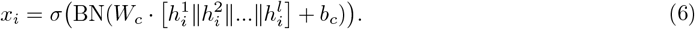

### Classifier and training

For prediction, the representation of candidate protein pair (*p*_*i*_, *p*_*j*_) is derived by element-wise multiplication of their embeddings, followed by a three-layer multilayer perceptron (MLP):

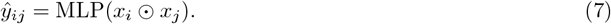

The task is formulated as a multi-label classification problem, in which each pair of proteins may participate in multiple types of PPIs. We utilize a multi-task binary cross-entropy loss for training, defined as:

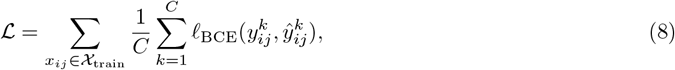

where *C* is the number of PPI types, 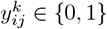 denotes the ground-truth label for type 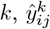 is the predicted probability, and ℓ_BCE_(*y, ŷ*) = −*y* log *ŷ* − (1 − *y*) log(1 − *ŷ*).

## Results

We benchmark DMG-PPI on SHS27K and SHS148K [30, 33, 38, 39], two canonical multi-type PPI datasets curated from STRING [40], a comprehensive resource for human interactome analysis. Both datasets were filtered to retain only high-confidence interactions (score ≥ 0.7), resulting in 12,517 PPIs in SHS27K and 44,488 PPIs in SHS148K. To minimize protein overlap between the training and test sets, we adopt the Breadth-First Search (BFS) and Depth-First Search (DFS) splitting strategies proposed by Lv et al. [39]. The benchmark methods include PIPR [41], ESM2 AMP [42], TUnA [43], GNN PPI [32], AFTGAN [33], LDMGNN [44], HIGH-PPI [30], BaPPI [34], and MAPE-PPI [31].

### DMG-PPI achieves the best performance and model capabilities

Figure 3a presents the results of DMG-PPI compared with leading benchmark methods. Each experiment was repeated five times, with mean values (two decimal places) reported; standard deviations are provided in S1 Appendix. Prediction performance is quantified by: (i) Micro-F1: the harmonic mean of precision and recall; (ii) AUPRC: area under the precision–recall curve; (iii) NDCG: normalized discounted cumulative gain; (iv) P@500L: proportion of positive PPIs among the top 500 ranked links; (v) AUROC: area under the receiver operating characteristic curve; and (vi) z-score aggregated across the above metrics.

**Fig 3:**
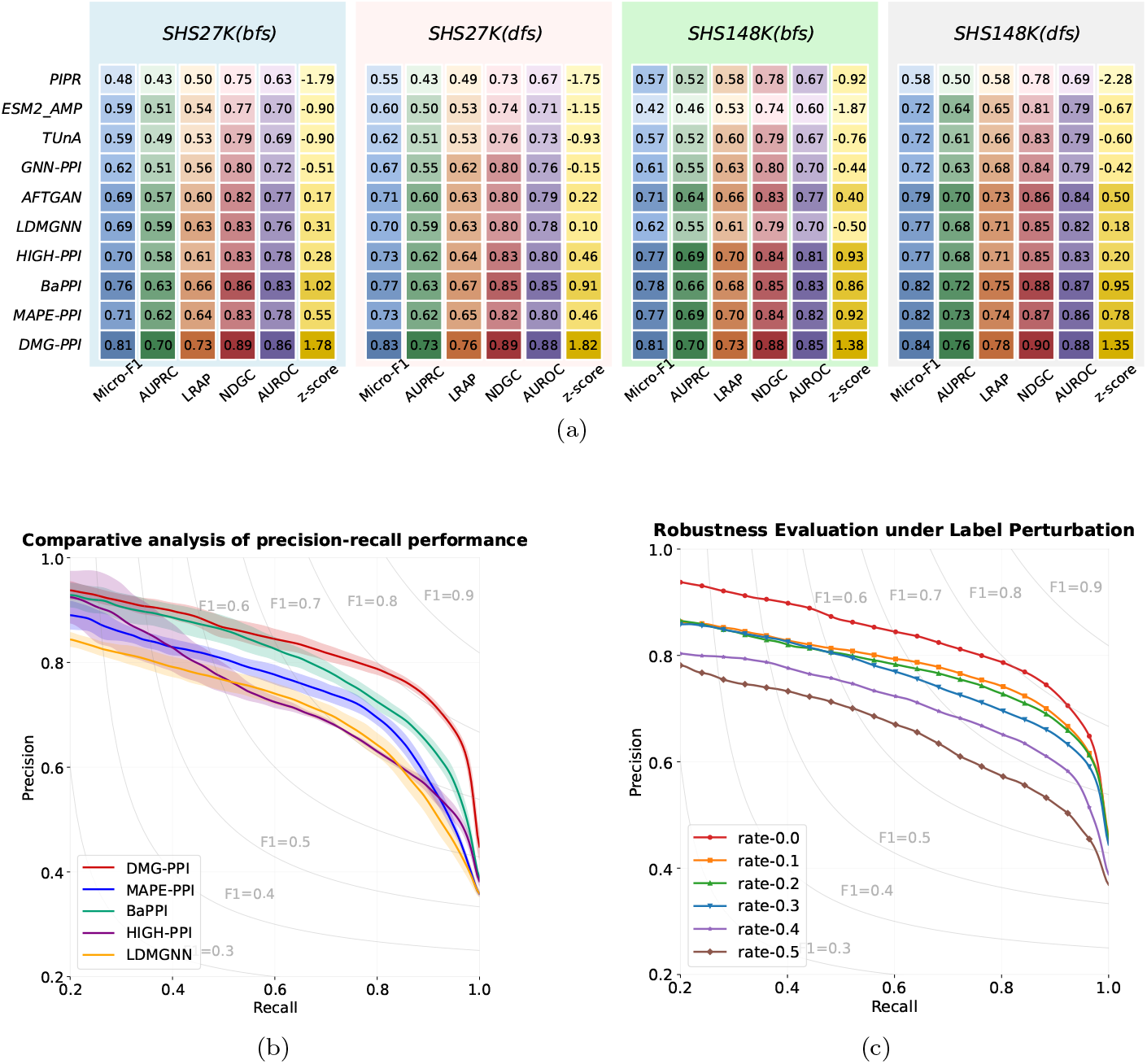
Comparative performance of DMG-PPI and benchmark methods. **(a)** Heatmaps showing the performance of each method across the following evaluation metrics: Micro-F1, AUPRC, LRAP, NDCG, AUROC, and overall z-score. Darker colors indicate better performance. **(b)** Precision–recall curves for PPI prediction on SHS27K, comparing the top five methods; shaded areas represent the range between the highest and lowest results. **(c)** Robustness of DMG-PPI to label perturbation on SHS27K, evaluated using precision–recall curves at different perturbation ratios

DMG-PPI consistently outperforms all state-of-the-art methods across every metric. MAPE-PPI and BaPPI achieve competitive second-place performance in different scenarios, while PIPR demonstrates notably inferior performance, likely due to its non-GNN architecture, which limits its ability to capture global topological information. Given the high class imbalance and skewed link distribution in PPI datasets, Micro-F1 provides a more reliable performance measure than NDCG or AUROC, as the latter tend to overestimate performance when generalization is weak [45]. This explains the smaller differences observed between methods under NDCG and AUROC. Quantitatively, DMG-PPI exceeds the second-best method in Micro-F1 by 7.19% on SHS27K and 3.14% on SHS148K.

We further evaluate robustness by introducing label perturbations into the training set to simulate undiscovered PPIs, which is a common feature of PPI datasets. Positive labels for each interaction were randomly removed at ratios ranging from 10–50% to simulate the frequently occurring undiscovered interactions. As shown in Fig 3c, DMG-PPI sustains an F1-score above 0.7 even with 30% label removal, with negligible performance loss when the ratio is below 30%. This resilience indicates that DMG-PPI captures stable global interaction patterns that persist despite incomplete supervision, particularly in high-recall regimes. At low recall, the model trained with complete labels retains a clearer advantage, underscoring robustness in top-ranked predictions.

Notably, the superior performance of DMG PPI can be attributed to its enhanced capability to recognize positive PPIs from the minority class. To illustrate this discrepancy, we examine four representative positive PPI cases formed by biologically important proteins. As shown in Fig 4, DMG PPI assigns systematically higher confidence scores to true positive interactions, with the minimum predicted probability exceeding 0.96. In contrast, the predicted probabilities produced by TuNA and HIGH-PPI for the same positive PPIs fall within a much lower range of 0.20 to 0.53, with most values concentrated between 0.3 and 0.4. DMG PPI exhibits a clear separation between positive and negative predictions, this pronounced separation in predicted probabilities highlights its superior discriminative power and reliability in identifying minority positive PPIs.

**Fig 4:**
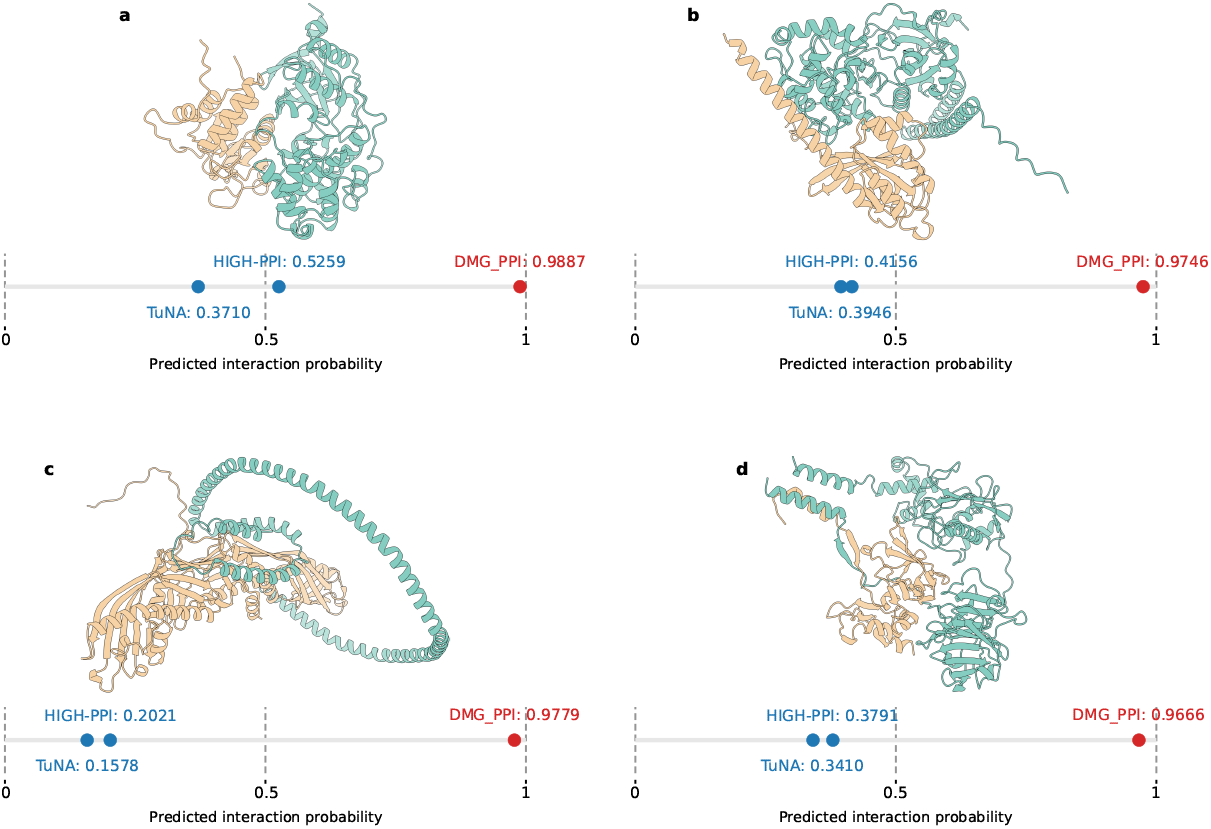
Protein–protein interaction (PPI) examples that are correctly predicted by DMG-PPI but not by HIGH-PPI or TuNA. The corresponding PPI complex structures are predicted using GRAMM and visualized with ChimeraX. **a** O15264 (Mitogen-activated protein kinase 13) and P51452 (Dual specificity protein phosphatase 3); **b** P11233 (Ras-related protein Ral-A) and Q15208 (Serine/threonine-protein kinase 38); **c** P11597 (Cholesteryl ester transfer protein) and P02647 (Apolipoprotein A-I); **d** P01033 (Metalloproteinase inhibitor 1) and Q9H306 (Matrix metalloproteinase-27).

### DMG-PPI precisely detects key interface residues underlying PPIs

Hotspots are specific subsets of residues that contribute disproportionately to the binding free energy, hence play a fundamental role in governing protein–protein interactions. The divergence message pathway (DMP) focuses on physicochemical and geometric complementarity at the protein interface, which are key factors underlying the formation of hotspots. The residue-level DMP scores allow estimation of the probability that a residue serves as such a site.

We systematically mask each residue individually and compute the resulting percentage change in both DMP and the prediction. A normalized Bayesian-style product of these two changes is then used to quantify the likelihood of each residue being a hotspot. We provide a case study of hotspot probability of interaction between HMBOX1 (UniProt ID O76528) and CSK (UniProt ID P41240). The proxy ground truth is computed by Rosetta Flex ddG. As shown in Fig 5a, the tendency of being hotspot is represented by color, DMG-PPI identifies nearly all hotspots with high accuracy. In particular, it successfully detects hotspots with low SASA (e.g., residue 296 of CSK, 3.76 Å^2^), which exhibit limited structural exposure and are often overlooked by conventional methods that rely solely on surface geometry. The ability of DMG-PPI to recognize these less apparent, structurally buried hotspots underscores its broader applicability and robustness in protein–protein interaction analysis.

**Fig 5:**
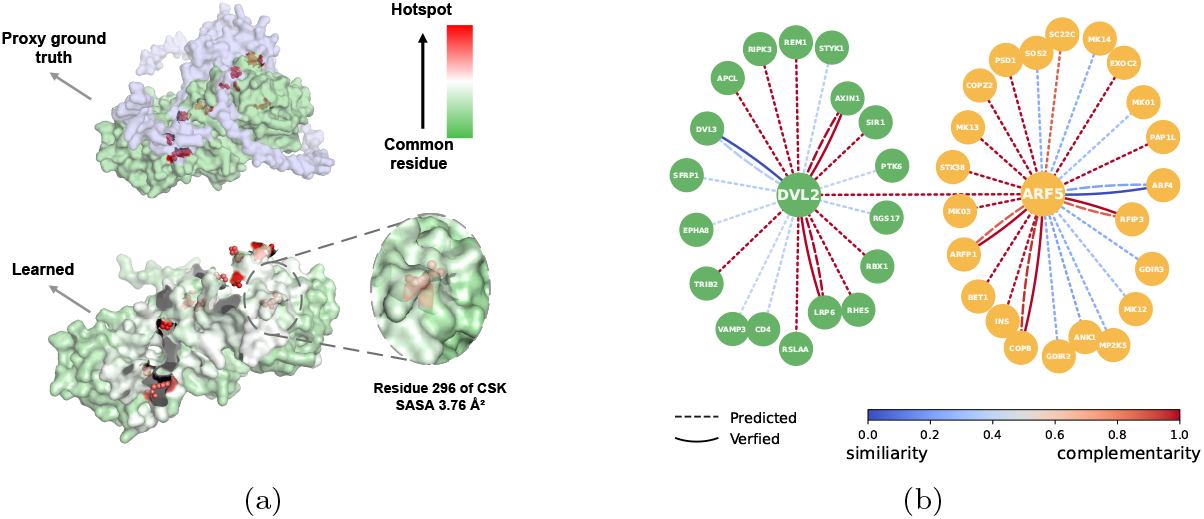
The interpretability provided by DMG-PPI. **(a)** Surface representation of a protein complex (UniProt IDs: O76528 and P41240). The likelihood of a residue being a hotspot is visualized using a color gradient from low (green) to high (red). **(b)** DMG-PPI accurately identify the interaction mode between proteins. Dashed lines represent the prediction computed by DMG-PPI, while solid lines represent ground-truth. The modes of PPIs are represented by edge color, varying along a spectrum from blue (similarity-driven) to red (complementarity-driven).

Furthermore, since the two channels of DMG-PPI correspond to shape complementarity and structural motifs, respectively, their associated gating coefficients can be leveraged to infer the dominant driving force of PPI. We provide a case study of the driven force that relates two proteins: DVL2, a well-connected, high-centrality node in Wnt signaling networks [46, 47], and ARF5, an essential functional protein [48] involved in vesicular trafficking, the ground truth is obtained from IntAct [49]. The confidence score of driven force is derived from the gating coefficients in our dual-channel framework. As shown in Fig 5b, experimentally verified driven force is represented by solid line, while prediction is indicated by dashed line, the color of edges demonstrate the confidence score of driven force. DMG-PPI correctly predicts the main driving force of all PPIs with available ground truth, with particularly strong accuracy observed for interactions such as DVL2–LPR6, DVL2–AXIN1, and ARF5–COPB. The accurate prediction provided by DMG-PPI can relieve the high cost of experimental method, the excellent interpreability also demonstrates DMG-PPI learns generalized protein representation.

### Learning dual interaction patterns enhances performance

The key innovation of our approach lies in a dual-channel framework that integrates distinct message-passing pathways, each tailored to a specific interaction pattern in the PPI network. To assess the contribution of learning dual interaction patterns, we compare the performance of the full dual-channel model with its single-channel variants, where only the AMP or DMP is used. The evaluation is conducted from the following perspectives: (1) an ablation study examining the individual contribution of each channel, (2) a type-specific performance analysis in terms of AUPR, and (3) an analysis of the interaction patterns among the top 500 predicted PPIs.

We first examine the performance of each channel on SHS27K and SHS148K. As illustrated in Fig 6a, the dual-channel framework substantially outperforms each individual message pathway on both datasets. These results indicate that each message pathway is limited to learning only a portion of the structural information inherent in the PPI network, thereby verifying the coexistence of interaction patterns and the effectiveness of our approach. Moreover, the variance of dual-channel framework is considerably lower than that of AMP and DMP, highlighting the model’s ability to achieve balanced learning across the two interaction patterns and its strong robustness. Notably, AMP performs better on SHS148K, whereas DMP yields more accurate predictions on SHS27K. This discrepancy may be attributed to the higher similarity among interacting proteins in SHS148K compared to SHS27K, which makes similarity-driven interactions more prominent in the former.

**Fig 6:**
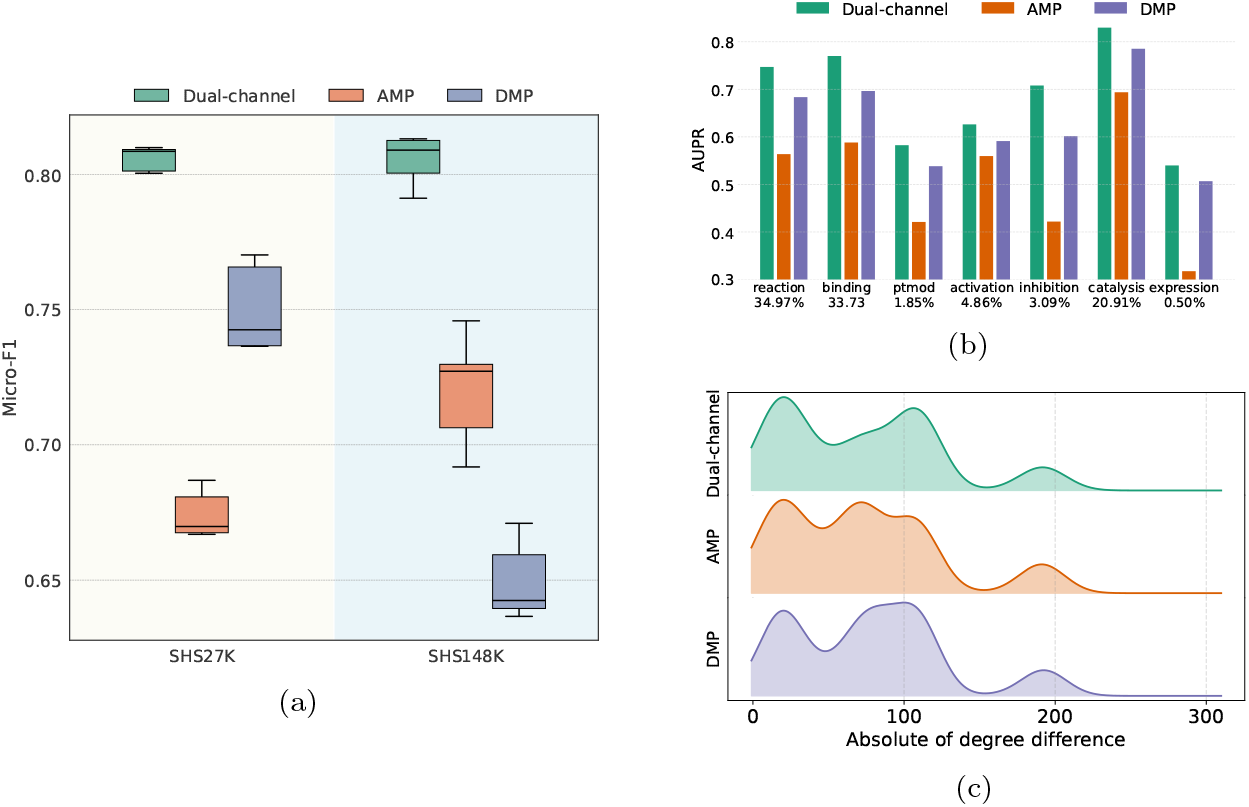
Performance comparison between each message pathway. **(a)** The Micro-F1 of each variant on SHS27K and SHS148K. **(b)** The AUPR across PPI types on SHS27K, the x-axis labels denote PPI types and the corresponding percentage. **(c)** Patterns of top-500 PPIs predicted by each variant. The x-axis represents the absolute value of the degree difference between the interacting proteins, while the y-axis indicates the percentage.

We further evaluate each channel’s performance across interaction types on the SHS27K dataset. In PPI networks, the distribution of interaction types is highly imbalanced: the dominant types (reaction, binding, and catalysis) account for over 80% of all interactions. Such skewed distribution may bias prediction models, making balanced performance across types a meaningful indicator of generalization. As shown in Fig 6b, the dual-channel framework significantly outperforms AMP and DMP across all interaction types. The performance gains of the dual-channel over AMP are especially pronounced for binding, ptmod, inhibition, and expression types that are strongly associated with biochemical complementarity (e.g., charge, shape, and hydrophobicity). These results demonstrate that our approach effectively addresses the gap in learning complementarity patterns within PPI networks. Particularly, ptmod (post-translational modification) interactions hold considerable biological significance, as they are central to protein function regulation, signal transduction pathways, and disease mechanisms. Improvements in predicting these interactions may thus have broad implications for downstream biomedical applications, including disease gene prioritization and drug target identification.

Finally, to probe the interaction patterns captured by each pathway, we analyze the top 500 predicted PPIs from the dual-channel, AMP, and DMP models on SHS27K. Specifically, we compare the absolute degree difference between interacting protein pairs. As illustrated in Fig 6c, certain recurring patterns emerge, such as peaks around degrees 25, 100, and 175. The distribution associated with the dual-channel model is more densely concentrated than those of AMP and DMP, suggesting that it captures more distinct structural characteristics and integrates them more effectively.

### DMG-PPI reveals the relationship between PPI types

In applications such as mechanistic modeling and pathway reconstruction [50], information about individual PPI types is often insufficient. Investigating the relationships between different PPI types can yield deeper mechanistic insights into molecular functions and disease mechanisms. To explore these inter-type functional dependencies, we decompose the classifier’s weight matrix into a set of *k × k* vectors based on the corresponding PPI type. The L2 norm of each vector is used to quantify the influence between PPI types.

The inferred influence map produced by DMG-PPI is presented in Fig 7, where each row indicates how all PPI types contribute to a specific PPI type. Notably, the inferred relationships align with some known biological mechanisms. For example, the reaction type is most influenced by ptmod and catalysis, consistent with previous findings that reaction events frequently occur in catalytic or modifying contexts [51, 52]. Overall, reaction, binding, ptmod, and catalysis types exhibit stronger dependence on other interaction types, suggesting a mechanistic interdependence that underlies their functional roles in molecular interaction events.

**Fig 7:**
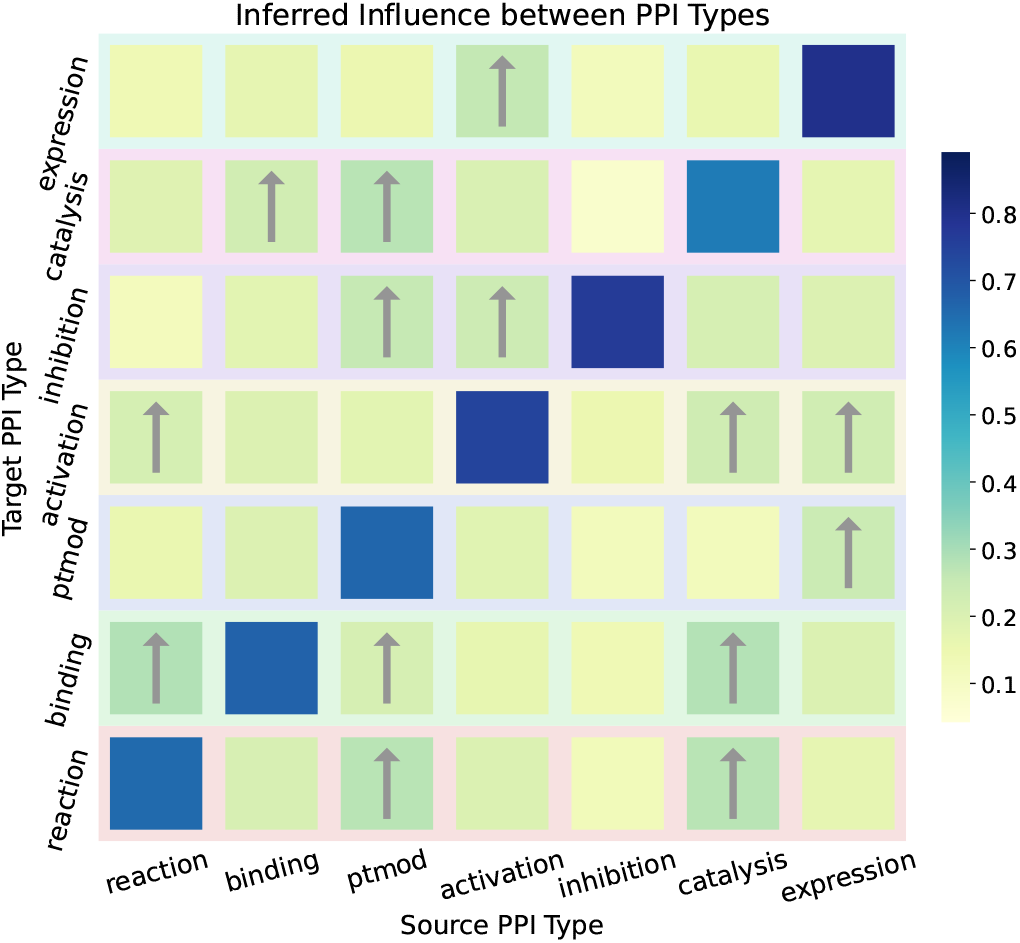
Directional influence among PPI types. The value at the *i*-th row and *j*-th column represents the influence of type *j* on type *i*, as inferred from the trained model. Influences with values greater than 0.2 are marked with arrows.

### DMG-PPI learns topology-robust representations beyond local motif density

To assess whether DMG-PPI learns unbiased representations that are not strongly influenced by local network topology, we compared embedding distance distributions for edges located in motif-dense versus motif-poor neighborhoods. The motif density of each edge was quantified as the sum of L3 motifs associated with its two endpoints. Edges in the bottom 20% and top 20% of motif density were defined as motif-poor and motif-richness, respectively. As shown in Fig 8, the resulting embedding distance distributions largely overlap, with only minor differences observed between the two groups. These findings suggest that DMG-PPI does not preferentially encode interactions in dense topological motifs, but instead learns representations that are relatively insensitive to local motif overrepresentation, consistent with a balanced integration of network context and protein-specific features.

**Fig 8:**
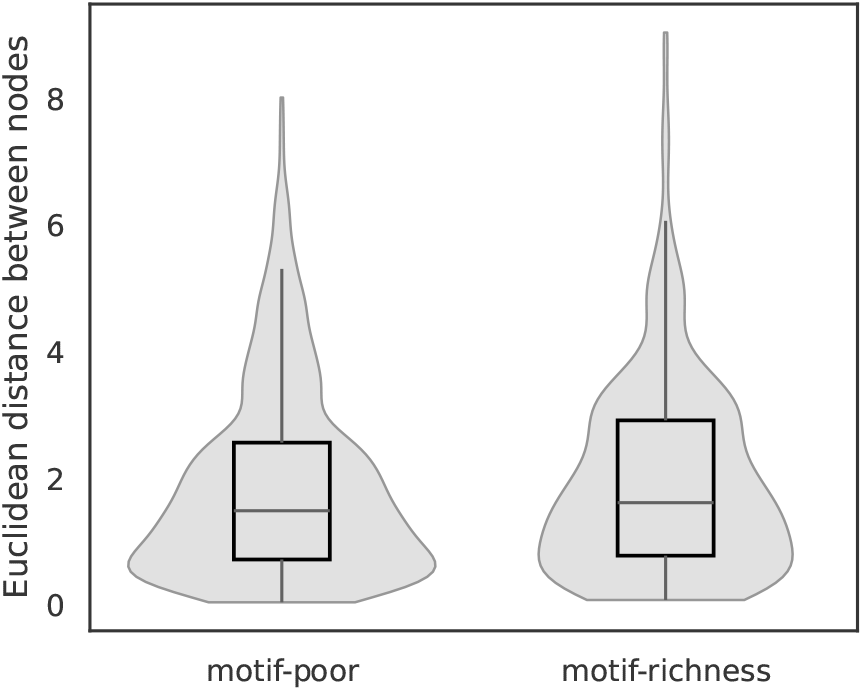
Distribution of Euclidean distances between node embeddings for edges in motif-richness versus motif-poor regions. The similarity of distances across regions suggests that DMG-PPI learns representations largely independent of local motif density.

Further, linear regression analysis indicates that motif richness explains only 1.6% of the variance in embedding distances (*R*^2^ = 0.016), suggesting that DMG-PPI embeddings are largely independent of local motif density. Consistently, Spearman correlation analysis reveals a weak monotonic relationship between motif richness and embedding distance (*ρ* = 0.10, *p* = 1.72 *×* 10^−8^), highlighting that the relative organization of edges in the embedding space is minimally influenced by motif structure. Together, these results demonstrate that DMG-PPI captures topology-robust representations that extend beyond the influence of local motif density.

## Discussion

In this paper, we introduce DMG-PPI, an efficient PPI prediction method that models the complex interaction patterns in PPI networks from a dual-channel GNN perspective. In the preprocessing stage, the physical connection and covalent bond are extracted as initial protein representation. At interactome level, we employ two channels that use additive and multuplicative aggregation to capture complementarity-driven and similarity-driven interactions, respectively. Experiments on classical benchmark datasets demonstrate that DMG-PPI significantly outperforms state-of-the-art methods in PPI prediction. Our dual-channel framework also exhibits strong generalization to minority PPI types and robustness under varying perturbation ratios. Moreover, given that surface complementarity is a primary determinant of binding specificity, DMG-PPI effectively identifies interaction modes and key residues of PPI, offering high interpretability. Overall, the dual-channel framework provides a powerful tool for systematic characterization of interactome connectivity and enhances mechanistic insights into PPIs.

While DMG-PPI achieves promising performance, some areas are worth consideration for further improvements. (1) This study primarily investigated the driving forces of PPIs in the context of binary interactions. However, in multiprotein complexes, the overall stability and functionality often emerge from a mosaic of interaction modes across different protein pairs. Modeling these interdependencies could provide a more faithful representation of the biophysical principles governing complex formation. (2) The potential of a unified protein graph that simultaneously incorporates physical and chemical features remains unexplored. A geometric GNN that directly operates on the full 3D atomic coordinates—integrating shape, electrostatics, and residue-level properties into a single graph could enable the learning of physically interpretable embeddings, potentially improving generalization to unseen proteins. (3) Protein language models can be leveraged to extract information about evolutionary patterns, sequence motifs, and functional signals.

Incorporating embeddings that implicitly encode multiple sequence alignment (MSA) information could be particularly helpful for predicting functional residues.

## Supporting information

## S1 Appendix

**Detailed experimental results on SHS27K with BFS scheme.**

S1 Appendix. Detailed experimental results on SHS27K with BFS scheme.
| method | Micro-F1 | AUPRC | LRAP | NDGC | AUROC | z-score |
| --- | --- | --- | --- | --- | --- | --- |
| DMG-PPI | $0.81 \pm 0.00$ | $0.70 \pm 0.01$ | $0.73 \pm 0.01$ | $0.89 \pm 0.00$ | $0.86 \pm 0.00$ | 1.56 |
| MAPE-PPI | $0.71 \pm 0.01$ | $0.62 \pm 0.01$ | $0.64 \pm 0.01$ | $0.83 \pm 0.01$ | $0.78 \pm 0.01$ | 0.32 |
| BaPPI | $0.76 \pm 0.01$ | $0.63 \pm 0.01$ | $0.66 \pm 0.01$ | $0.87 \pm 0.01$ | $0.83 \pm 0.01$ | 0.80 |
| HIGH-PPI | $0.70 \pm 0.00$ | $0.58 \pm 0.00$ | $0.61 \pm 0.01$ | $0.83 \pm 0.01$ | $0.78 \pm 0.00$ | 0.04 |
| LDMGNN | $0.69 \pm 0.02$ | $0.59 \pm 0.01$ | $0.63 \pm 0.01$ | $0.83 \pm 0.01$ | $0.76 \pm 0.01$ | 0.09 |
| AFTGAN | $0.69 \pm 0.01$ | $0.57 \pm 0.02$ | $0.60 \pm 0.01$ | $0.83 \pm 0.01$ | $0.77 \pm 0.00$ | -0.04 |
| GNN-PPI | $0.62 \pm 0.02$ | $0.51 \pm 0.02$ | $0.56 \pm 0.02$ | $0.81 \pm 0.01$ | $0.72 \pm 0.01$ | -0.74 |
| TU<sub>n</sub>A | $0.59 \pm 0.02$ | $0.49 \pm 0.01$ | $0.53 \pm 0.02$ | $0.79 \pm 0.01$ | $0.69 \pm 0.01$ | -0.90 |
| ESM2_AMP | $0.59 \pm 0.03$ | $0.51 \pm 0.01$ | $0.54 \pm 0.02$ | $0.77 \pm 0.01$ | $0.70 \pm 0.01$ | -0.90 |
| PIPR | $0.48 \pm 0.05$ | $0.43 \pm 0.02$ | $0.50 \pm 0.01$ | $0.75 \pm 0.01$ | $0.63 \pm 0.02$ | -2.03 |

**Detailed experimental results on SHS27K with DFS scheme**.

| method | Micro-F1 | AUPRC | LRAP | NDGC | AUROC | z-score |
| --- | --- | --- | --- | --- | --- | --- |
| DMG-PPI | $0.83 \pm 0.01$ | $0.73 \pm 0.01$ | $0.76 \pm 0.00$ | $0.89 \pm 0.01$ | $0.88 \pm 0.01$ | 1.65 |
| MAPE-PPI | $0.73 \pm 0.00$ | $0.62 \pm 0.00$ | $0.65 \pm 0.00$ | $0.81 \pm 0.00$ | $0.80 \pm 0.00$ | 0.18 |
| BaPPI | $0.77 \pm 0.01$ | $0.63 \pm 0.01$ | $0.67 \pm 0.01$ | $0.85 \pm 0.01$ | $0.85 \pm 0.00$ | 0.69 |
| HIGH-PPI | $0.73 \pm 0.00$ | $0.62 \pm 0.01$ | $0.64 \pm 0.02$ | $0.83 \pm 0.01$ | $0.80 \pm 0.00$ | 0.21 |
| LDMGNN | $0.70 \pm 0.01$ | $0.59 \pm 0.01$ | $0.63 \pm 0.01$ | $0.80 \pm 0.00$ | $0.78 \pm 0.01$ | -0.17 |
| AFTGAN | $0.71 \pm 0.01$ | $0.60 \pm 0.01$ | $0.63 \pm 0.02$ | $0.81 \pm 0.02$ | $0.79 \pm 0.01$ | -0.03 |
| GNN-PPI | $0.67 \pm 0.01$ | $0.55 \pm 0.01$ | $0.62 \pm 0.01$ | $0.80 \pm 0.01$ | $0.76 \pm 0.00$ | -0.44 |
| TUnA | $0.62 \pm 0.01$ | $0.51 \pm 0.01$ | $0.53 \pm 0.01$ | $0.76 \pm 0.01$ | $0.73 \pm 0.01$ | -0.93 |
| ESM2_AMP | $0.60 \pm 0.01$ | $0.50 \pm 0.01$ | $0.53 \pm 0.01$ | $0.74 \pm 0.01$ | $0.71 \pm 0.02$ | -1.15 |
| PIPR | $0.55 \pm 0.02$ | $0.43 \pm 0.01$ | $0.49 \pm 0.01$ | $0.73 \pm 0.01$ | $0.67 \pm 0.01$ | -2.10 |

**Detailed experimental results on SHS148K with BFS scheme**.

| method | Micro-F1 | AUPRC | LRAP | NDGC | AUROC | z-score |
| --- | --- | --- | --- | --- | --- | --- |
| DMG-PPI | $0.81 \pm 0.01$ | $0.70 \pm 0.01$ | $0.73 \pm 0.01$ | $0.88 \pm 0.01$ | $0.85 \pm 0.01$ | 1.32 |
| MAPE-PPI | $0.77 \pm 0.00$ | $0.69 \pm 0.01$ | $0.70 \pm 0.01$ | $0.85 \pm 0.01$ | $0.82 \pm 0.00$ | 0.75 |
| BaPPI | $0.78 \pm 0.02$ | $0.66 \pm 0.03$ | $0.68 \pm 0.04$ | $0.85 \pm 0.02$ | $0.83 \pm 0.01$ | 0.68 |
| HIGH-PPI | $0.77 \pm 0.00$ | $0.69 \pm 0.01$ | $0.70 \pm 0.01$ | $0.85 \pm 0.01$ | $0.81 \pm 0.00$ | 0.76 |
| LDMGNN | $0.62 \pm 0.00$ | $0.55 \pm 0.02$ | $0.61 \pm 0.03$ | $0.79 \pm 0.01$ | $0.70 \pm 0.01$ | -1.04 |
| AFTGAN | $0.71 \pm 0.01$ | $0.64 \pm 0.01$ | $0.66 \pm 0.01$ | $0.83 \pm 0.00$ | $0.77 \pm 0.00$ | 0.09 |
| GNN-PPI | $0.61 \pm 0.01$ | $0.55 \pm 0.01$ | $0.63 \pm 0.01$ | $0.80 \pm 0.00$ | $0.70 \pm 0.00$ | -0.98 |
| TUnA | $0.57 \pm 0.01$ | $0.52 \pm 0.01$ | $0.60 \pm 0.01$ | $0.79 \pm 0.01$ | $0.67 \pm 0.01$ | -0.76 |
| ESM2_AMP | $0.42 \pm 0.01$ | $0.46 \pm 0.01$ | $0.53 \pm 0.01$ | $0.74 \pm 0.02$ | $0.60 \pm 0.01$ | -1.87 |
| PIPR | $0.57 \pm 0.01$ | $0.52 \pm 0.02$ | $0.58 \pm 0.02$ | $0.78 \pm 0.01$ | $0.67 \pm 0.01$ | -1.57 |

**Detailed experimental results on SHS148K with DFS scheme**.

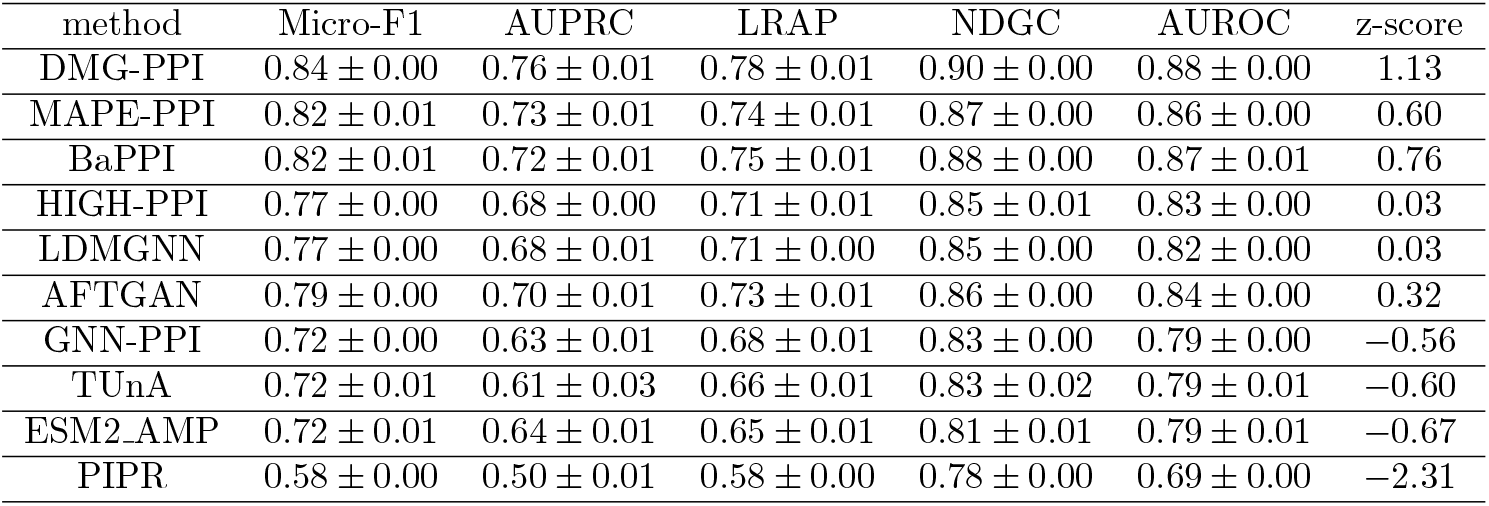

| method | Micro-F1 | AUPRC | LRAP | NDGC | AUROC | z-score |
| --- | --- | --- | --- | --- | --- | --- |
| DMG-PPI | $0.84 \pm 0.00$ | $0.76 \pm 0.01$ | $0.78 \pm 0.01$ | $0.90 \pm 0.00$ | $0.88 \pm 0.00$ | 1.13 |
| MAPE-PPI | $0.82 \pm 0.01$ | $0.73 \pm 0.01$ | $0.74 \pm 0.01$ | $0.87 \pm 0.00$ | $0.86 \pm 0.00$ | 0.60 |
| BaPPI | $0.82 \pm 0.01$ | $0.72 \pm 0.01$ | $0.75 \pm 0.01$ | $0.88 \pm 0.00$ | $0.87 \pm 0.01$ | 0.76 |
| HIGH-PPI | $0.77 \pm 0.00$ | $0.68 \pm 0.00$ | $0.71 \pm 0.01$ | $0.85 \pm 0.01$ | $0.83 \pm 0.00$ | 0.03 |
| LDMGNN | $0.77 \pm 0.00$ | $0.68 \pm 0.01$ | $0.71 \pm 0.00$ | $0.85 \pm 0.00$ | $0.82 \pm 0.00$ | 0.03 |
| AFTGAN | $0.79 \pm 0.00$ | $0.70 \pm 0.01$ | $0.73 \pm 0.01$ | $0.86 \pm 0.00$ | $0.84 \pm 0.00$ | 0.32 |
| GNN-PPI | $0.72 \pm 0.00$ | $0.63 \pm 0.01$ | $0.68 \pm 0.01$ | $0.83 \pm 0.00$ | $0.79 \pm 0.00$ | -0.56 |
| TUnA | $0.72 \pm 0.01$ | $0.61 \pm 0.03$ | $0.66 \pm 0.01$ | $0.83 \pm 0.02$ | $0.79 \pm 0.01$ | -0.60 |
| ESM2_AMP | $0.72 \pm 0.01$ | $0.64 \pm 0.01$ | $0.65 \pm 0.01$ | $0.81 \pm 0.01$ | $0.79 \pm 0.01$ | -0.67 |
| PIPR | $0.58 \pm 0.00$ | $0.50 \pm 0.01$ | $0.58 \pm 0.00$ | $0.78 \pm 0.00$ | $0.69 \pm 0.00$ | -2.31 |

## Data availability statement

All synthetic data and code necessary to reproduce the results of this study are publicly available in the github repository https://github.com/ttan6729/DMG_PPI and google drive link.

